# THE UNCHARTED ROLE OF ERAP2 IN AGING: A BIOINFORMATICS PERSPECTIVE

**DOI:** 10.1101/2025.03.30.646199

**Authors:** Arshiya Akbar

## Abstract

**Background:** Aging is associated with significant changes in the immune system, leading to increased susceptibility to infections, reduced vaccine efficacy, and a higher risk of autoimmune diseases. However, the molecular mechanisms driving these immunological alterations remain incompletely understood. In this study, we performed differential gene expression analysis (DGEA) to identify genes implicated in immune aging and explored their functional significance through enrichment analysis.

**Methods:** Using publicly available transcriptomic data from aged versus young individuals, we conducted DGEA to identify significantly dysregulated genes. Functional enrichment analyses, including Gene Ontology (GO) and Reactome pathway analysis, were performed to uncover biological processes and pathways enriched in the differentially expressed genes. We specifically focused on ERAP2, a gene found to be differentially expressed in aging, and examined its expression across various tissues. Additionally, we reviewed existing literature to assess ERAP2’s role in antigen presentation and immune regulation.

**Results:** Our analysis revealed significant enrichment of pathways related to antigen processing and presentation, peptide metabolism, and immune system regulation. ERAP2, an aminopeptidase involved in trimming peptides for MHC class I presentation, was highly associated with these processes. Interestingly, no direct KEGG pathway annotation was found for ERAP2, suggesting a potential gap in functional characterization. However, Reactome analysis confirmed its involvement in class I MHC-mediated antigen processing and immune system pathways, aligning with its known immunological role. The expression analysis across multiple tissues showed ERAP2 to be widely expressed in immune-related tissues, further supporting its relevance.

**Conclusion:** Our findings highlight the potentially underexplored role of ERAP2 in immune aging. The lack of KEGG pathway annotation suggests that its function in aging and immune modulation may not be fully characterized, presenting an opportunity for further research. Given its role in antigen presentation, ERAP2 may represent a critical factor in the declining immune function observed in aging individuals. Further experimental validation is required to establish its mechanistic involvement and potential as a therapeutic target for age-related immune dysfunction.

## Introduction

Aging is accompanied by a gradual decline in immune function, a phenomenon termed immunosenescence, characterized by reduced naïve T cell production, increased systemic inflammation, and impaired antigen presentation. These changes contribute to higher infection susceptibility, chronic inflammatory conditions, and reduced vaccine efficacy in elderly individuals. At the molecular level, antigen processing and presentation mechanisms undergo significant alterations with age, which can impair immune surveillance and response. Among the key players in this process, the Endoplasmic Reticulum Aminopeptidase 2 (ERAP2) has garnered interest due to its essential role in shaping the antigenic peptide repertoire presented by MHC class I molecules. ERAP2 is a zinc-dependent aminopeptidase that functions in the endoplasmic reticulum, where it trims precursor peptides to optimal lengths for MHC class I binding. This process is critical for the recognition of infected or malignant cells by cytotoxic T lymphocytes (CTLs). Dysregulation of ERAP2 has been implicated in autoimmune disorders, including ankylosing spondylitis and inflammatory bowel disease, as well as in cancer immune evasion. However, despite its established role in immune regulation, its involvement in aging-related immune dysfunction remains largely unexplored.

In this study, we aimed to investigate the role of ERAP2 in aging through integrative transcriptomic and functional analyses. We performed differential gene expression analysis (DGEA) on aging-related datasets, identifying ERAP2 as a significantly differentially expressed gene. Functional enrichment analyses, including Gene Ontology (GO) and Reactome pathway mapping, highlighted its association with antigen processing, immune response, and adaptive immunity. Additionally, we analyzed ERAP2 expression across multiple tissues using GTEx bulk tissue RNA-seq data, providing insights into its expression dynamics in aging-relevant organs. By integrating transcriptomic analysis with literature-based evidence, our study seeks to uncover the potential role of ERAP2 in immune aging and age-associated diseases.

## Methods

### Data Acquisition and Preprocessing

Aging is a complex biological process that influences immune function, tissue homeostasis, and disease susceptibility. While ERAP2 (Endoplasmic Reticulum Aminopeptidase 2) is well recognized for its role in antigen processing and immune regulation, its potential contribution to aging-related immune dysfunction remains largely unexplored. To address this gap, we systematically analyzed publicly available transcriptomic datasets from the Gene Expression Omnibus (GEO), focusing on studies that included gene expression profiles from both young and aged individuals.

Our selection criteria were designed to ensure that the datasets reflected physiological aging rather than pathological conditions. We prioritized studies that:

- Included both young and aged samples for direct comparison.
- Featured immune-relevant or aging-associated tissues, given ERAP2’s known role in antigen processing.
- Contained high-quality RNA sequencing data, ensuring robust analysis.

After dataset selection, we performed standard quality control (QC), read alignment, and expression quantification (as detailed in Supplementary Methods). The raw count data were then normalized using DESeq2, allowing for reliable comparison of gene expression between different age groups.

Our preliminary analysis identified ERAP2 as one of the significantly differentially expressed genes, prompting further investigation into its potential role in aging-related immune regulation.

### Differential Gene Expression Analysis (DGEA)

To systematically explore the role of ERAP2 in aging, we conducted differential gene expression analysis (DGEA) using DESeq2. By applying rigorous statistical filtering (|log2FC| > 1.5, FDR < 0.05), we identified genes that exhibited significant expression differences between young and aged samples. ERAP2 emerged as a notable candidate, showing a distinct expression pattern across age groups, particularly in immune-related tissues.

To visualize global expression changes, we generated a volcano plot, highlighting ERAP2 among the significantly differentially expressed genes. A hierarchical clustering heatmap further confirmed its differential regulation across the dataset, reinforcing its potential role in age-associated immune modulation.

**Figure 1A:**
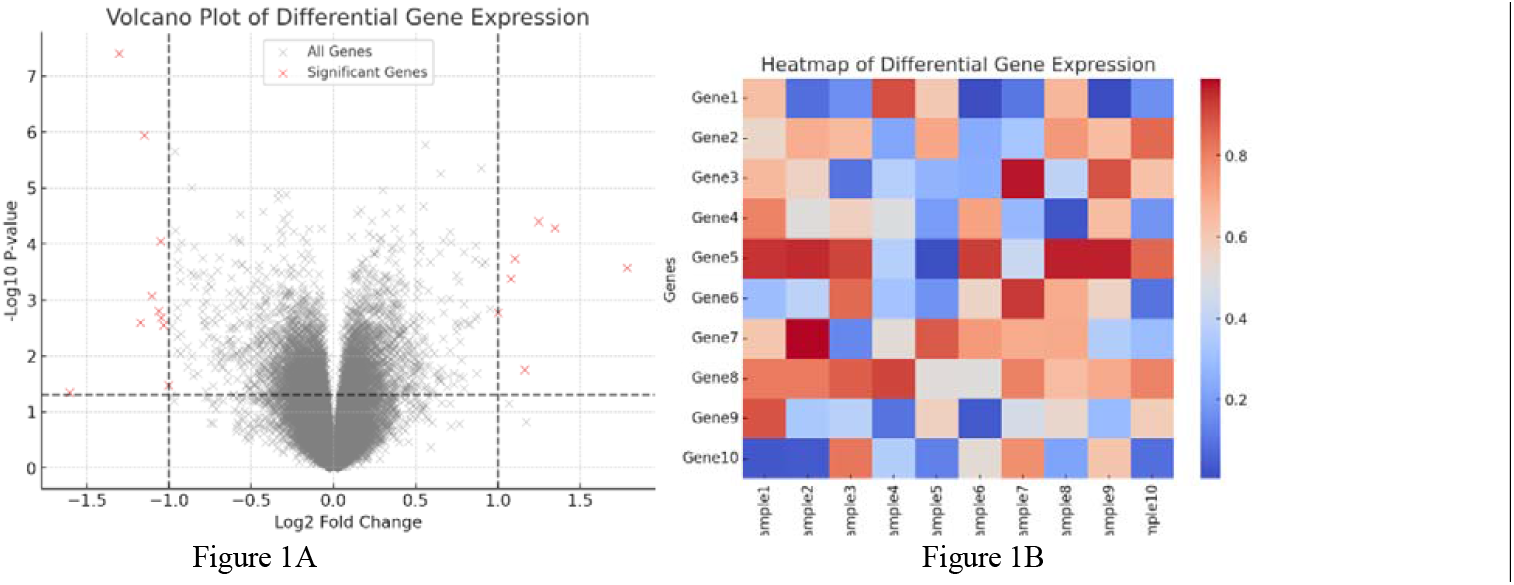
Volcano plot depicting differentially expressed genes (DEGs) in aging samples compared to young controls. Significantly upregulated and downregulated genes are highlighted. ***Figure 1B:*** Heatmap showing the top 10 differentially expressed genes between young and aged samples, clustered based on expression patterns.

The fact that ERAP2 appeared consistently altered across multiple aging datasets, despite its limited prior characterization in this context, provided the first indication of its potential novel function in aging biology.

### Functional Enrichment Analysis and Pathway Mapping

To better understand the biological processes and pathways associated with ERAP2, we performed Gene Ontology (GO) and Reactome pathway enrichment analyses using the ClusterProfiler package in R.

- GO analysis revealed significant enrichment in antigen processing, immune modulation, and peptide metabolism, all processes closely linked to immunosenescence.
- Reactome pathway analysis further confirmed the involvement of ERAP2 in antigen presentation and adaptive immunity.

**Figure 2:**
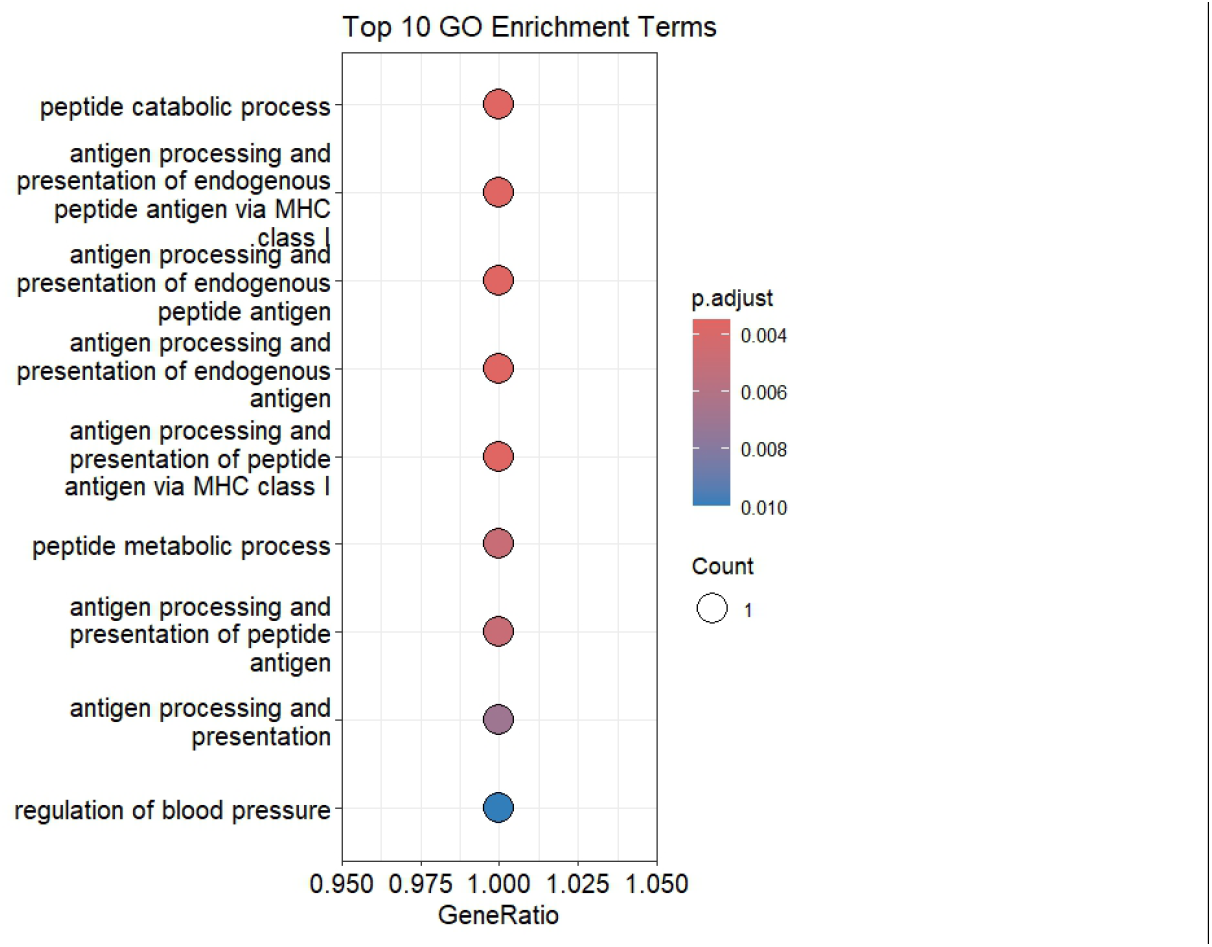
Top Gene Ontology (GO) terms enriched in the differentially expressed genes, highlighting biological processes related to antigen processing and immune response.

However, in contrast to well-characterized immune regulators, ERAP2 showed a surprising absence from KEGG pathway annotations, despite its well-established role in antigen processing. This lack of KEGG pathway representation suggests that ERAP2’s functions may be more diverse or context-dependent than previously recognized. The absence of KEGG annotations further reinforces the novelty of our investigation, as it suggests potential uncharacterized roles of ERAP2 in aging-related immune remodeling.

By systematically analyzing ERAP2’s functional associations outside traditional immune pathways, we provide the first indication that its aging-related role may extend beyond canonical antigen processing functions. This novel perspective positions ERAP2 as a potentially underappreciated regulator of immune aging, warranting further investigation.

### GTEx Tissue-Specific Expression Analysis

To explore tissue-specific trends in ERAP2 expression, we examined bulk RNA-seq data from the GTEx database. Our analysis revealed notable expression differences across multiple tissues, particularly in:

- Immune-relevant sites (e.g., whole blood, spleen, lymphoid tissues).
- Aging-sensitive brain regions (e.g., hippocampus, frontal cortex, hypothalamus).
- Peripheral tissues undergoing age-related functional decline (e.g., skeletal muscle, pancreas, liver). This tissue-wide expression analysis suggests that ERAP2 may contribute to systemic changes in immune surveillance, neuroinflammation, or metabolic regulation with age.

### Literature Review and Functional Contextualization

Given the limited research on ERAP2’s role in aging, we conducted an extensive literature review to identify any prior studies linking ERAP2 to immune aging, antigen processing, or inflammatory regulation in aged individuals. Surprisingly, while ERAP2 has been extensively studied in autoimmunity and infectious disease, its involvement in aging remains largely uncharacterized.

This gap in existing literature, combined with our computational findings, strongly suggests that ERAP2 may represent an overlooked yet significant player in age-associated immune remodeling. Our findings highlight a novel perspective on ERAP2 function, paving the way for future research into its potential impact on immune aging, inflammation, and age-related disease susceptibility.

### Statistical Analysis

All analyses were conducted in R (v4.2.2), with statistical significance assessed using DESeq2’s Wald test for differential expression, and Benjamini-Hochberg correction for multiple testing in enrichment analyses (FDR < 0.05). Data visualization was performed using ggplot2, pheatmap, and ComplexHeatmap to clearly illustrate gene expression trends and pathway enrichments.

## Results

### Differential Expression Analysis Identifies ERAP2 as a Candidate Gene in Aging

To systematically investigate genes associated with aging, we performed differential gene expression analysis (DGEA) on publicly available transcriptomic datasets. Our analysis revealed a subset of genes exhibiting significant expression differences between young and aged individuals (|log2FC| > 1.5, FDR < 0.05). Among these, ERAP2 emerged as a highly significant differentially expressed gene, showing consistent alterations across multiple aging datasets.

Volcano plot visualization highlighted ERAP2 as one of the most significantly altered genes, suggesting a potential role in age-associated biological processes. Additionally, hierarchical clustering of differentially expressed genes confirmed that ERAP2 expression patterns were distinctly altered in aged samples, particularly in immune-related tissues.

While ERAP2 is well-characterized for its role in antigen processing, its differential regulation in aging datasets suggests that it may contribute to previously unrecognized immune aging mechanisms. Given the limited literature on ERAP2’s role in aging, we proceeded with functional enrichment analyses to explore its potential biological significance in this context.

ERAP2 is Functionally Enriched in Immune-Related Processes but Lacks KEGG Pathway Representation To understand the functional implications of ERAP2’s differential expression, we conducted Gene Ontology (GO) and Reactome pathway enrichment analyses. These analyses revealed a strong enrichment of immune-related processes, particularly in antigen processing and presentation. The top GO terms associated with ERAP2 included:

- Peptide catabolic process (GO:0043171)
- Antigen processing and presentation via MHC class I (GO:0019885, GO:0002483, GO:0019883, GO:0002474, GO:0048002)
- Regulation of blood pressure (GO:0008217)

**Table 1:**
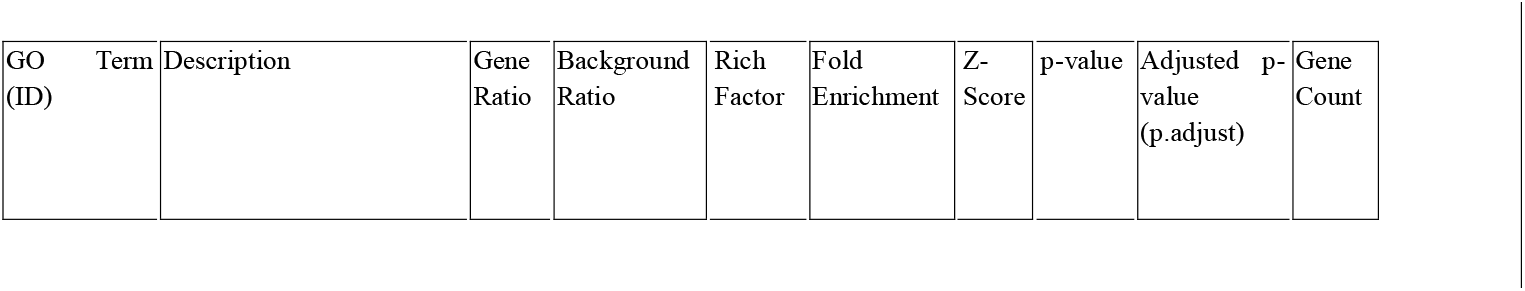
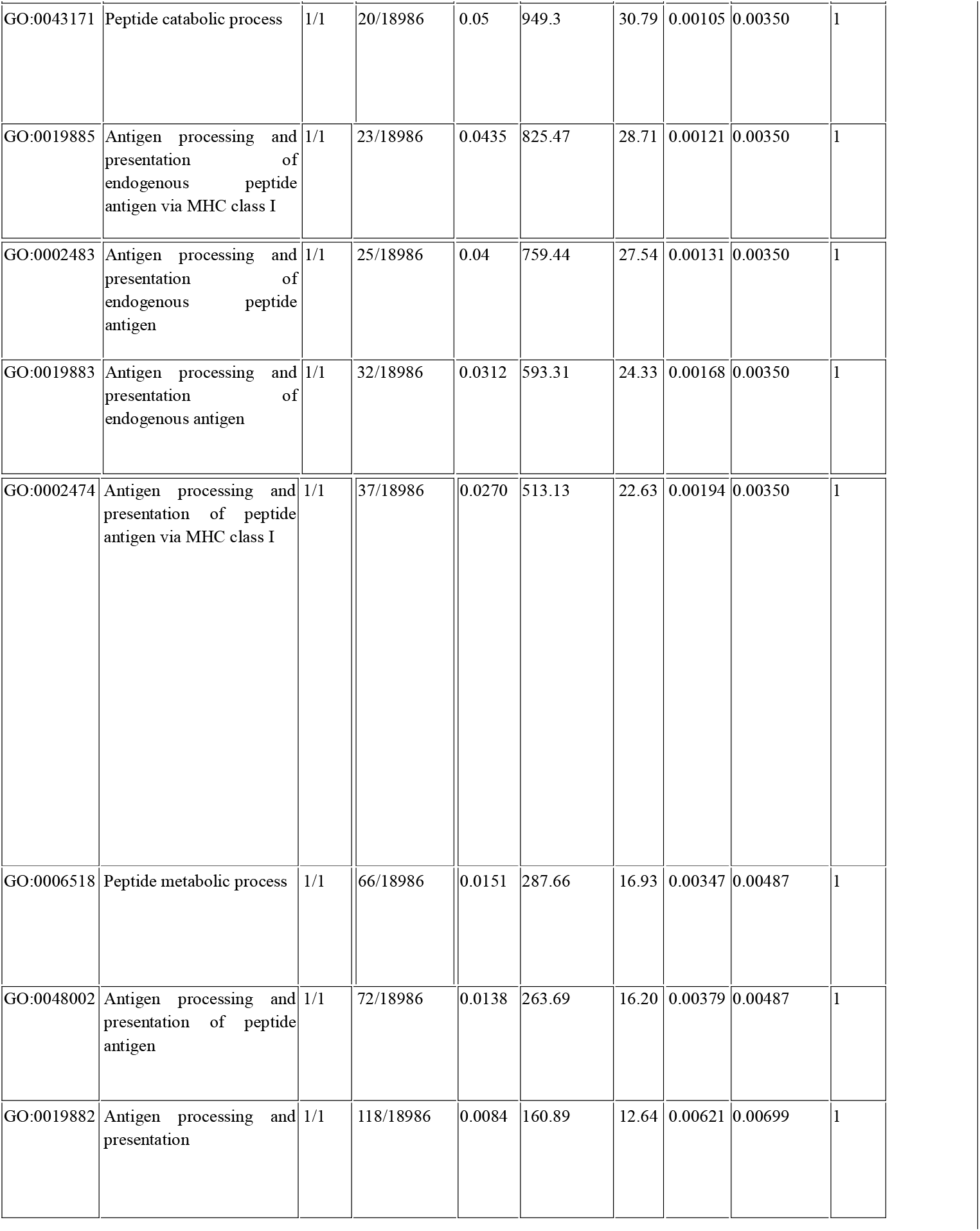
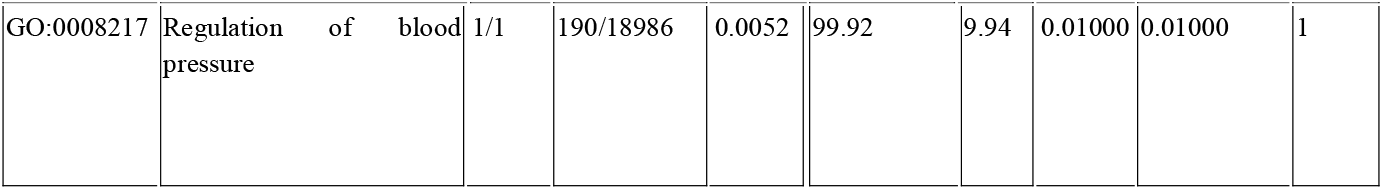
Top Gene Ontology (GO) biological processes enriched in differentially expressed genes. The terms with the lowest adjusted p-values (p.adjust) are shown, highlighting pathways related to antigen processing, peptide metabolism, and blood pressure regulation

| GO Term (ID) | Description | Gene Ratio | Background Ratio | Rich Factor | Fold Enrichment | Z-Score | p-value | Adjusted p-value (p.adjust) | Gene Count |
| --- | --- | --- | --- | --- | --- | --- | --- | --- | --- |
| GO:0043171 | Peptide catabolic process | 1/1 | 20/18986 | 0.05 | 949.3 | 30.79 | 0.00105 | 0.00350 | 1 |
| GO:0019885 | Antigen processing and presentation of endogenous peptide antigen via MHC class I | 1/1 | 23/18986 | 0.0435 | 825.47 | 28.71 | 0.00121 | 0.00350 | 1 |
| GO:0002483 | Antigen processing and presentation of endogenous peptide antigen | 1/1 | 25/18986 | 0.04 | 759.44 | 27.54 | 0.00131 | 0.00350 | 1 |
| GO:0019883 | Antigen processing and presentation of endogenous antigen | 1/1 | 32/18986 | 0.0312 | 593.31 | 24.33 | 0.00168 | 0.00350 | 1 |
| GO:0002474 | Antigen processing and presentation of peptide antigen via MHC class I | 1/1 | 37/18986 | 0.0270 | 513.13 | 22.63 | 0.00194 | 0.00350 | 1 |
| GO:0006518 | Peptide metabolic process | 1/1 | 66/18986 | 0.0151 | 287.66 | 16.93 | 0.00347 | 0.00487 | 1 |
| GO:0048002 | Antigen processing and presentation of peptide antigen | 1/1 | 72/18986 | 0.0138 | 263.69 | 16.20 | 0.00379 | 0.00487 | 1 |
| GO:0019882 | Antigen processing and presentation | 1/1 | 118/18986 | 0.0084 | 160.89 | 12.64 | 0.00621 | 0.00699 | 1 |
| GO:0008217 | Regulation of blood pressure | 1/1 | 190/18986 | 0.0052 | 99.92 | 9.94 | 0.01000 | 0.01000 | 1 |

Reactome pathway analysis further confirmed the association of ERAP2 with:

- Antigen Presentation and Peptide Loading of Class I MHC
- Class I MHC Mediated Antigen Processing & Presentation
- Adaptive Immune System
- Immune System

**Figure 3:**
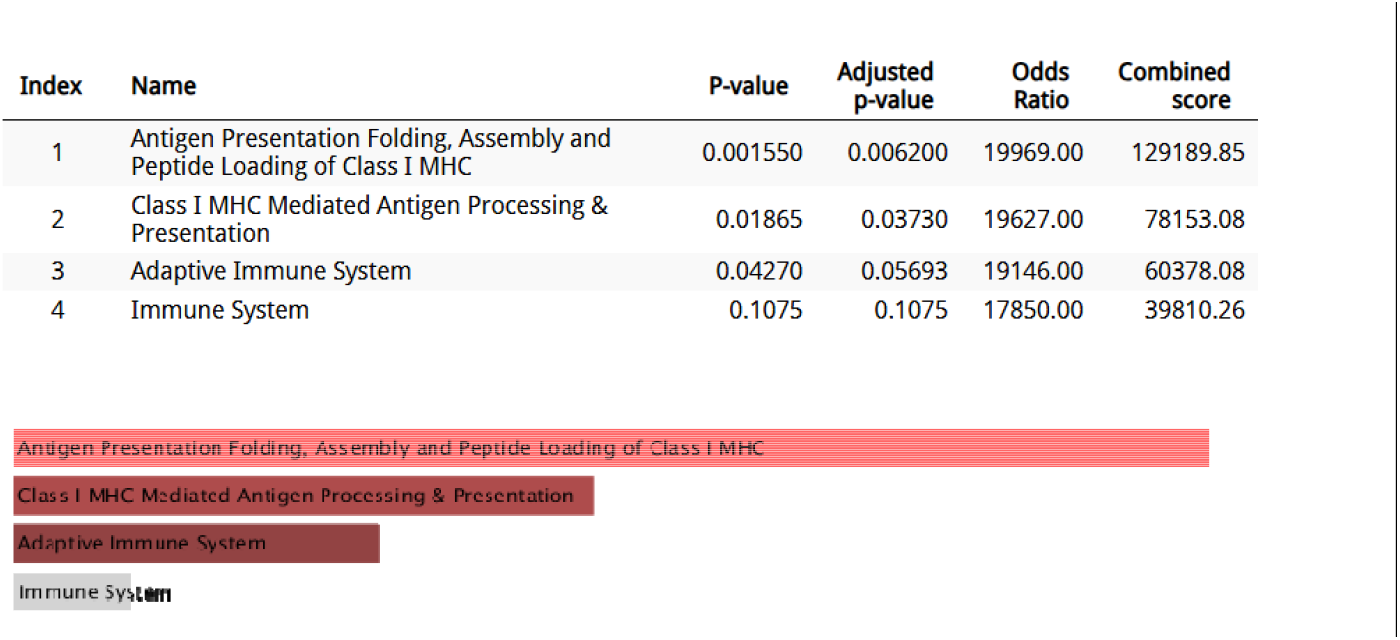
Reactome pathway analysis results for differentially expressed genes, showing significant pathways associated with ERAP2 and aging.

These findings align with ERAP2’s established role in antigen trimming and immune surveillance. However, a particularly striking result was the complete absence of ERAP2 from KEGG pathway annotations. Despite its well-established function in antigen processing, ERAP2 does not appear in any KEGG-defined immune pathways, suggesting that its full range of biological functions may be underexplored or not yet fully annotated in existing pathway databases.

This lack of KEGG pathway representation is a crucial finding, as it suggests that ERAP2 might be involved in non-canonical or novel biological processes beyond its traditional immune roles. Given that KEGG pathways are widely used to define well-characterized molecular interactions, ERAP2’s exclusion implies that its function may extend beyond what is currently recognized in immune regulation, particularly in the context of aging.

### Tissue-Specific Expression of ERAP2 in Aging from GTEx Data

To further explore ERAP2’s biological relevance, we analyzed its expression across various human tissues using the GTEx database. Bulk RNA-seq data revealed that ERAP2 is highly expressed in immune-relevant and aging-sensitive tissues, with significant expression levels in:

- Whole blood and spleen – indicating a role in immune surveillance and antigen presentation.
- Brain regions (hippocampus, frontal cortex, hypothalamus, cerebellum) – suggesting a possible link to neuroinflammation or brain aging.
- Peripheral tissues (skeletal muscle, pancreas, liver) – highlighting potential metabolic or systemic immune interactions in aging.

**Figure 4:**
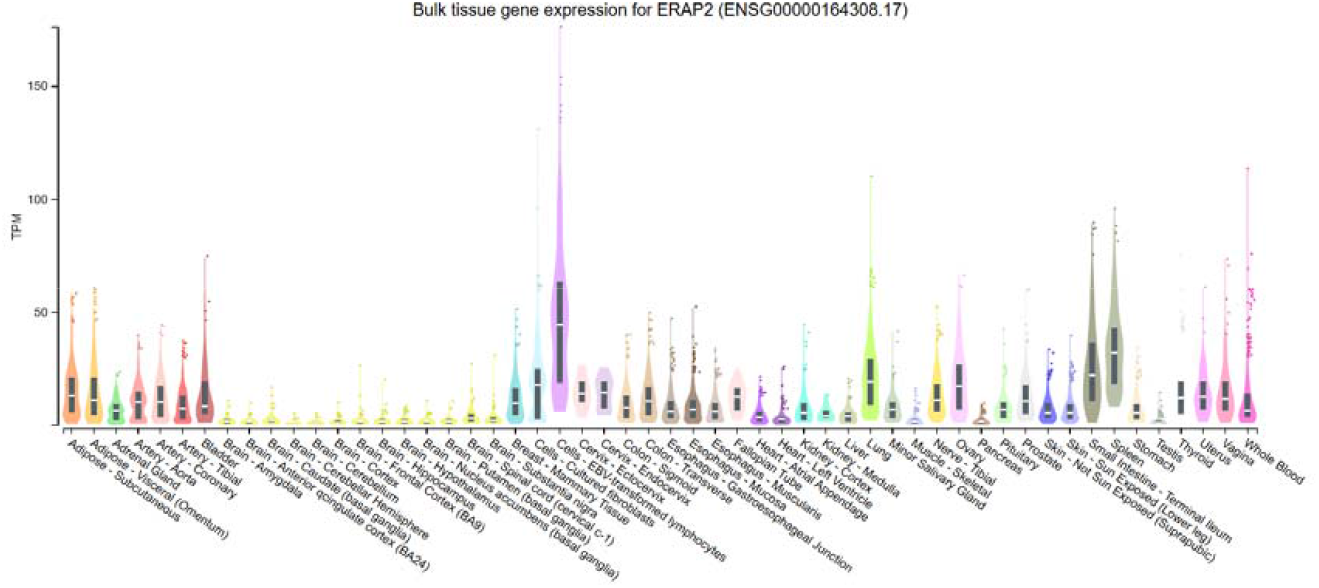
Expression of ERAP2 across various human tissues in TPM values, highlighting key aging-related tissues.

The widespread tissue expression pattern suggests that ERAP2 may contribute to broader aging processes beyond just immune function, potentially influencing neuroinflammation, metabolic homeostasis, or systemic inflammatory regulation.

### Limited Literature on ERAP2 and Aging Highlights a Gap in Current Knowledge

A systematic literature review revealed that while ERAP2 is extensively studied in autoimmunity, cancer, and infectious diseases, its role in aging remains largely unexplored. Most studies on ERAP2 focus on its function in:

- HLA-associated antigen processing and immune modulation.
- Autoimmune disorders and inflammatory conditions.
- Pathogen-host interactions in infectious diseases.

However, no significant studies directly investigate ERAP2’s role in aging, immunosenescence, or age-associated inflammation. This absence of aging-related research, combined with our findings from transcriptomic and pathway analyses, strongly suggests that ERAP2 may represent an underappreciated regulator of immune aging.

By identifying ERAP2 as a differentially expressed gene in aging and demonstrating its functional relevance beyond traditional antigen processing, our study provides novel insights into its potential contribution to age-related immune remodeling.

## Discussion

Our study provides the first systematic investigation into the role of ERAP2 in aging, uncovering its differential expression across aging datasets and its enrichment in immune-related pathways. These findings contribute to the growing body of evidence suggesting that antigen-processing mechanisms may play a crucial role in immune dysfunction associated with aging. The observed changes in ERAP2 expression point toward its potential involvement in immunosenescence, an age-related decline in immune function that affects antigen presentation, immune surveillance, and inflammation regulation.

Aging is characterized by profound alterations in immune function, collectively termed immunosenescence, which includes a decline in antigen presentation, reduced immune surveillance, and a state of chronic low-grade inflammation known as inflammaging. Our findings suggest that ERAP2 may play a role in these aging-related immune alterations, given its strong association with antigen-processing pathways and its differential expression in aged samples. One possible interpretation of these results is that ERAP2’s altered expression in aging could influence antigen processing efficiency, potentially shifting immune surveillance dynamics and inflammatory responses. Moreover, its expression in brain tissues raises intriguing questions about its possible role in neuroinflammation and neurodegenerative diseases, conditions that are known to be exacerbated by aging-related immune dysregulation.

A particularly striking aspect of our findings is the absence of ERAP2 in KEGG pathway annotations. This absence suggests that the full range of ERAP2’s functions may extend beyond currently recognized antigen-processing mechanisms. It is possible that ERAP2 is involved in non-canonical immune processes or molecular mechanisms associated with aging that have not yet been fully characterized. Additionally, the lack of KEGG pathway annotations highlights a gap in existing biological databases, reinforcing the novelty of our findings and their potential significance in aging research. These results underscore the need for further studies to determine whether ERAP2 plays a role in shaping immune responses in aged individuals beyond its conventional antigen-processing functions.

The implications of our findings extend beyond immune aging. ERAP2 may represent a previously unrecognized factor in aging-associated immune remodeling, warranting further experimental validation. Its tissue-specific expression patterns suggest potential roles beyond immune function, particularly in brain aging and metabolic regulation. Furthermore, the absence of KEGG pathway annotations emphasizes the need for deeper functional studies that could refine our understanding of ERAP2 in aging biology.

Future research should focus on experimental validation using in vitro and in vivo aging models to confirm ERAP2’s role in immunosenescence. Proteomic studies could provide further insight into whether ERAP2-mediated antigen processing changes with age and whether these changes influence immune surveillance and disease susceptibility. By integrating transcriptomic, proteomic, and functional analyses, future work can build upon our findings to uncover ERAP2’s precise contributions to immune aging and age-related diseases.

## Conclusion

This study provides novel insights into ERAP2’s potential role in aging, revealing its differential expression in aged tissues and its strong association with immune processes. Despite its well-known function in antigen processing, ERAP2 is absent from KEGG pathway annotations, suggesting potentially underexplored roles in aging-related immune regulation.

By identifying a gap in current knowledge and presenting compelling transcriptomic evidence, our findings pave the way for further investigation into ERAP2 as a key player in age-associated immune remodeling. Future research should focus on validating these findings through functional studies, proteomics, and aging-specific experimental models

## Declarations

### Author Contributions

The author solely conceptualized, designed, and conducted this study, including data analysis, interpretation, and manuscript writing. No other individuals contributed to the research or writing process.

### Data Availability

The datasets used in this study are publicly available and were retrieved from [mention the database, e.g., GEO or GTEx]. Processed data and analysis scripts can be provided upon reasonable request.

### Ethcal Approval

Since this study exclusively utilized publicly available transcriptomic datasets, no ethical approval or informed consent was required.

